# Comparing utility functions between risky and riskless choice in rhesus monkeys

**DOI:** 10.1101/2021.01.12.426382

**Authors:** Philipe M. Bujold, Simone Ferrari-Toniolo, Leo Chi U Seak, Wolfram Schultz

## Abstract

Decisions can be risky or riskless, depending on the outcomes of the choice. Expected Utility Theory describes risky choices as a utility maximization process: we choose the option with the highest subjective value (utility), which we compute considering both the option’s value and its associated risk. According to the random utility maximization framework, riskless choices could also be based on a utility measure. Neuronal mechanisms of utility-based choice may thus be common to both risky and riskless choices. This assumption would require the existence of a utility function that accounts for both risky and riskless decisions. Here, we investigated whether the choice behavior of macaque monkeys in riskless and risky decisions could be described by a common underlying utility function. We found that the utility functions elicited in the two choice scenarios were different from each other, even after taking into account the contribution of subjective probability weighting. Our results suggest that distinct utility representations exist for riskless and risky choices, which could reflect distinct neuronal representations of the utility quantities, or distinct brain mechanisms for risky and riskless choices. The different utility functions should be taken into account in neuronal investigations of utility-based choice.

## Introduction

Whether we are choosing between fruits or vegetables at the supermarket, deciding to jaywalk in the face of incoming traffic, or picking the ideal friends to go traveling with, most of our decisions fall under two categories: some have certain outcomes, some do not. Economists call these risky or riskless decisions (‘risk’ referring to the uncertainty of a choice’s outcome), and-while vastly untested – there is general agreement in economics that peoples’ preferences in one type of situation parallels preferences in the other.

In economics, Expected Utility Theory (EUT) (von Neumann and Morgenstern, 1944) served as the dominant model of risky decision-making until the inception of behavioral economics in the 1970s. Under EUT, a decision-maker’s attitude towards risk was fully captured by the curvature of their utility function: a mapping of reward quantities onto an internal, subjective metric. A concave utility function predicted an aversion to risk, while a convex one predicted risk-seeking behavior. Importantly, EUT assumed that the utility of a riskless choice option could be computed through the same utility function used for risky options. On the other hand, experimental findings indicated discrepancies between risky and riskless utility functions (Barron et al., 1984; Stalmeier and Bezembinder, 1999).

Contrasting with EUT, Prospect Theory (PT) highlighted a difference between risky and riskless choices through the introduction of subjective probability weightings. Rather than being solely predicted by an individual’s utility curvature, one’s risk-attitude would also vary with their subjective treatment of outcome probabilities (Kahneman and Tversky, 1979; Tversky and Kahneman, 1992). In other words, while EUT assumed that risk attitudes derived exclusively from the way in which people value rewards, PT made the case for two components: the curvature of the utility function and the subjective weighting of probability.

PT has since become widespread in the study of risky and riskless decision-making (Kahneman et al., 1990; Lattimore et al., 1992; Camerer et al., 2002; Hertwig and Erev, 2009). With all the studies on behavior that make use of PT, there is a remarkable lack of research validating its predictions in both risky and riskless choices; the limitation being that risky utilities (or PT values) are usually measured from choices between risky options (Stott, 2006; Tversky & Kahneman, 1992) while this clearly cannot be done in a riskless context. One interesting avenue has been to compare risky and riskless preferences via introspective metrics. In a study by Stalmeier & Bezembinder (1999), medical patients were asked questions that involved risky outcomes:”would you rather: live 20 years with a migraine on x days per week (followed by death), or live 20 years with a p% chance of getting migraines y times a week, z times a week otherwise”; and questions where all options were riskless:”which difference is larger: the difference between 0 days of migraine and x days of migraine, or the difference between x days of migraine and 3 days of migraine”. Modelling preferences through PT, they found that risky and riskless utilities were identical, and that probability weighting accounted for most of the discrepancy between the risk attitudes predicted by riskless utilities and the risk attitudes measured from risky choices. A similar approach by Abdellaoui, Barrios, & Wakker (2007), this time using money outcomes (gains) rather than medical outcomes (losses), led to the similar conclusion: PT successfully reconciled risky and riskless utilities.

Since the subjects in these studies were generally risk-averse (for gains), it remains to be seen whether PT also reconciles risky and riskless utilities for risk-seeking decision-makers. Additionally, the results of these introspective studies have recently been challenged by a set of studies using a more modern, incentive-compatible approach: the use of time trade-offs as means to study riskless decisions (Cheung, 2016). In these studies, people make choices between larger rewards delivered in the future (with certainty) and smaller rewards delivered now; utilities from intertemporal choices are then compared to those estimated from risky choices. Unlike introspective experiments, however, the majority of the research done on time trade-offs reports discrepancies between riskless, time-discounted utility functions and risky ones (Musallam et al., 2004; Andreoni and Sprenger, 2012; Abdellaoui et al., 2013; Cheung and L., 2015; Lopez-Guzman et al., 2018), but see Andersen et al., 2008)- discrepancies that even probability weighting cannot resolve.

The lack of clear insight as to PT’s ability to reconcile risky and riskless choices represents a crucial limitation to the interpretation of this fundamental economic model; particularly as it rapidly became the de facto model of choice to study animal behavior and neuroeconomics (De Martino et al., 2006; Lakshminarayanan et al., 2011; Marshall and Kirkpatrick, 2013; Stauffer et al., 2015; Chen and Stuphorn, 2018; Farashahi et al., 2018; Ferrari-Toniolo et al., 2019a). Simultaneously, since there have been no attempts at reconciling risky and riskless utilities in nonhuman decision-makers, there is no evidence to suggest that either human interpretations can be used to explain animals’ choice behavior.

The present study explores the link between the risky and riskless utilities of our close primate relative: the rhesus macaque. We presented monkeys with two types of binary choice trials: risky trials, where monkeys made choices between certain and uncertain juice rewards, and riskless trials, which only included choices between two certain juice magnitudes. We elicited the shape of the utility curves in the two domains, using the random utility maximization (RUM) framework (for review, see McFadden, 2001) in combination with a PT-based discrete choice model. Importantly, this risky/riskless design addressed two of the most important caveats in human studies: (i) both risky and riskless trials were incentive compatible (relying on revealed preferences rather than introspection), and (ii) choices were presented in the exact same way for both risky and riskless decisions.

By parametrically separating the contributions that utility and weighted probability had on the monkeys’ risky choices, we found that, just like the human studies had previously shown, risky utilities were closer to riskless utilities once probability weighting had been accounted for. We did not, however, find that these utilities were identical, suggesting that two different utility quantities or mechanisms could drive behavior in risky and riskless choices.

## Methods

### Animals

Two male rhesus macaques (*Macaca mulatta*; Monkey A: 11.2 kg, Monkey B: 15.3 kg) participated in this experiment. All animals used in the study were born in captivity, at the Medical Research Council’s Centre for Macaques (CFM) in the UK. The animals were pair-housed for most of the experiment and had previous experience with the visual stimuli and experimental setup (Ferrari-Toniolo et al., 2019). The animals repeatedly chose between two reward options (reward-predicting stimuli) presented on an upright computer monitor. While sitting in a primate chair (Crist instruments), they used a left-right joystick (Biotronix Workshop, the University of Cambridge) to indicate their choice on each trial and received the reward they selected at the end of each of these binary choice trials (Fig. 1a).

**Figure 1.**
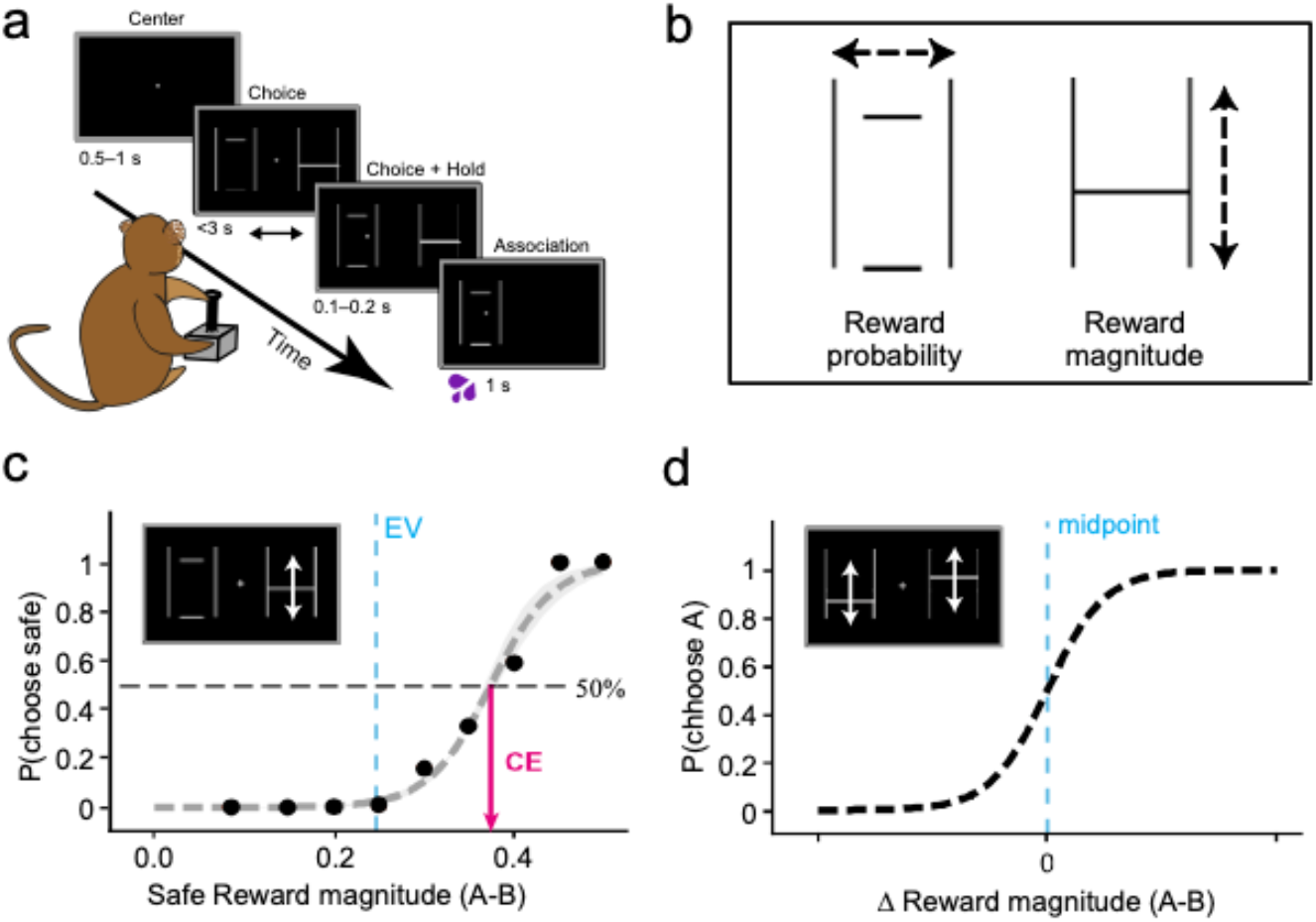
Experimental design and measures of risky and riskless choices. a) **Binary choice task.** The monkeys chose one of two gambles with a left-right motion joystick. They received the blackcurrant juice reward associated with the chosen stimuli after each trial. Time, in seconds, indicate the duration of each of the task’s main events. b) **Schema of visual stimuli.** Rewards were visually represented by horizontal lines (one or two) set between two vertical ones. The vertical position of these lines signalled the magnitude of said rewards. The width of these lines, the probability that these rewards would be realized). c) **Estimating certainty equivalents from risky choices.** Monkeys chose between a safe reward and a risky gamble on each trial. The safe rewards alternated pseudorandomly on every trial – they could be of any magnitude between 0 ml and 0.5 ml in 0.05 ml increments. Each point is a measure of choice ratio: the monkey’s probability of choosing the gamble option over various safe rewards. Psychometric softmax functions (Eq. 1) were fit to these choice ratios, then used to measure the certainty equivalents (CEs) of individual gambles (the safe magnitude for which the probability of either choice was 0.5; black arrow). The solid vertical line indicates the expected value (EV) of the gamble represented in the box. d) **Estimating the strength of preferences from riskless choices.** Riskless safe rewards were presented against one another, the probability of choosing the higher magnitude option (A) is plotted on the y-axis as a function of the difference in magnitude between the two options presented (Δ magnitude). The differences in magnitude tested were 0.02 ml, 0.04 ml, 0.06 ml, and a psychometric curve, anchored with its inflection anchored at a Δ magnitude of 0, were fit on the choice ratios measured (Eq. 2). These functions were fit to different magnitude levels, and the temperature of each curve was linked to the strength of monkeys’ preferences at each of these different levels.

This research has been ethically reviewed, approved, regulated, and supervised by the following institutions and individuals in the UK and at the University of Cambridge (UCam): the Minister of State at the UK Home Office, the Animals in Science Regulation Unit (ASRU) of the UK Home Office implementing the Animals (Scientific Procedures) Act 1986 with Amendment Regulations 2012, the UK Animals in Science Committee (ASC), the local UK Home Office Inspector, the UK National Centre for Replacement, Refinement and Reduction of Animal Experiments (NC3Rs), the UCam Animal Welfare and Ethical Review Body (AWERB), the UCam Governance and Strategy Committee, the Home Office Establishment License Holder of the UCam Biomedical Service (UBS), the UBS Director for Governance and Welfare, the UBS Named Information and Compliance Support Officer, the UBS Named Veterinary Surgeon (NVS), and the UBS Named Animal Care and Welfare Officer (NACWO).

### Task design and setup

The premise of this study was to compare the utility functions estimated from monkeys’ choices in risky or riskless decisions. To do so, monkeys were presented with sets of choices that could then be translated into utility metrics. The utilities measured from riskless choices were compared with utilities derived from risky choices, first assuming no subjective weighing of probabilities (EUT utilities), then accounting for the contribution of probability weighting (PT utilities).

Reward options took the form of various combinations of reward magnitude and probability, and were represented on the monitor through horizontal lines that scaled, and moved, relative to two vertical ‘framing’ lines (fig 1b). Reward magnitudes were represented by the vertical position of the horizontal lines: 0 ml at the bottom of the vertical frame (1.5ml at the top, and 0 < m < 1.5 in-between), whilst the probability of receiving said reward was represented by the width of the horizontal lines within the frame. A single, horizontal line that touched the frames at both ends signaled a certain reward (probability p = 1); multiple lines that failed to touch the frames indicated gambles with probabilistic outcomes, each with associated probability 0 < p < 1 (Fig. 1a). The monkeys were trained to associate these two-dimensional visual stimuli with blackcurrant juice rewards over the course of two years, and both monkeys had previous experience with the task and stimuli before this study. They had both experienced reward probabilities that ranged from 0 to 1 (Ferrari-Toniolo et al., 2019b), and reward magnitudes that ranged from 0 ml to 1.3 ml of juice. For this study, reward magnitudes were held between 0 ml and 0.5 ml of blackcurrant juice, and gamble options all had a probability of 0.5.

Each binary choice trial began with a white cross at the center of a black screen, if the monkey were holding the joystick, a cursor would also appear on the screen (Fig. 1a). Using the joystick, the monkeys initiated each trial by moving the cursor to the center cross and holding it there for 0.5-1s. Following this holding period, two reward options appeared to the left and to the right of the central cross (see Fig. 1a). The animal had 3s to convey his decision by moving the joystick to the selected side and holding his choice for 0.1-0.2s-the unselected option would then disappear. The selected option lingered on the screen for 1 s after reward delivery – followed by a variable inter-trial period of 1–2 s before the next trial. Errors were defined as unsuccessful central holds, side selection holds, or trials where no choices were made. Each of these resulted in a 6 s timeout for the animal, after which the trial would be repeated (ensuring the elicitation of preferences for each tested option pair). Additionally, all reward options were repeated on both the left and right sides of the computer screen, alternating pseudorandomly to control for any side preference. Both the joystick position and task event times were sampled and stored at 1 kHz on a Windows 7 computer running custom MATLAB software (The MathWorks, 2015a; Psychtoolbox version 3.0.11). we collected on average 423 ± 91 (SD) trials per session over 22 sessions for monkey A, and 338 ± 41 trials over 7 sessions for monkey B. Only trials where the option pair had been repeated at least 4 time were analyzed in this study. Data processing and statistical analyses were run in python (Python 3.7.3, SciPy 1.2.1, see Oliphant, 2007).

### Revealing preferences for risky and riskless choice

The monkeys’ daily reward preferences were measured in risky and riskless choice sequences under the framework of utility maximization. In risky choice sequences, trials always pit a risky gamble against a safe option – the utility of different reward magnitudes was estimated via the ratio of choices between different gamble and safe rewards. All of the gambles comprised two equally likely reward outcomes (though one could be 0 ml). In riskless choice sequences, monkeys were presented with pairs of ‘safe’ options with a single fixed outcome – we used the ratio of choice between pairs of rewards to estimate utility.

### Estimating utility functions in risky choice

For risky sequences, utilities were estimated using the fractile-bisection procedure – a method that involves dividing the range of possible utilities into progressively smaller halves (or fractals) and estimating the reward magnitude associated with each of these utility fractals. Simply put, the procedure defined set utility metrics (in this case ½, ¼ and ¾, and 1/8 and 7/8 of the maximum utility, see Fig. 2a, b) for which the corresponding safe rewards were derived (Fig. 2a).

**Figure 2.**
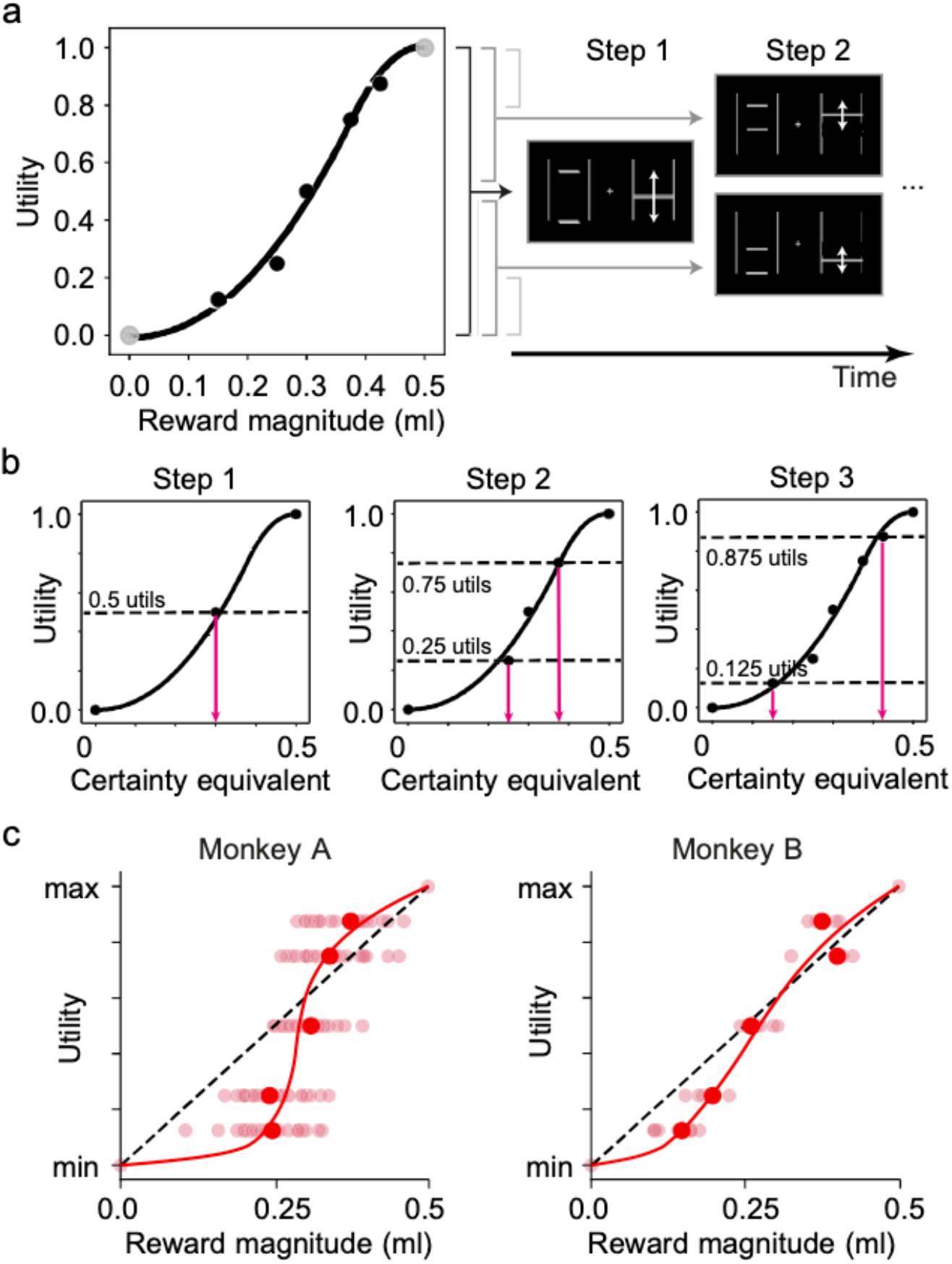
Estimating risky utilities using the fractile procedure. a) **Fixed utilities are mapped onto different reward magnitudes.** The gambles that monkeys experienced are defined from bisections of the range of possible reward magnitudes. For each step the gambles were held fixed; safe magnitudes varied by 0.05ml increments. b) **Estimation of utility using the stepwise, fractile method.** In step 1, the monkeys were presented with an equivariant gamble comprised of the maximum and minimum magnitudes in the tested reward range. The CE of the gamble was estimated and assigned a utility of 50%. In step 2, two new equivariant gambles were defined from the CE elicited in step 1. The CEs of these gambles were elicited and assigned a utility of 25% and 75%. Two more gambles are defined in step 3, from the CEs elicited in step 2. Their CEs were then assigned a utility of 12.5% and 87.5%. Parametric utility functions, anchored at 0 and 1, were fitted on these utility estimates (see methods). c) **Utility functions estimated from choices.** Datapoints represent daily CEs (semi-transparent) and their median values (red filled circles) tied to specific utility levels, as estimated through the fractile procedure. Both monkeys exhibit risk-seeking behaviour for low-magnitude rewards, and risk-aversion for high-magnitude ones. The data represents individual utility estimates gathered over 22 sessions for monkey A, and 7 sessions for monkey B. The red curves were obtained by fitting piecewise polynomial functions to the measured CEs (cubic splines with three knots).

Utility values of 0 and 1 were arbitrarily assigned to 0ml and 0.5ml of juice, respectively. Since monkeys only experienced trials set between these reward magnitudes, this constrained all utility estimates between a 0 and 1. Then, in accordance with EUT, a utility of 0.5 was assigned to the equiprobable gamble formed of these two magnitudes (0.5 = [0.5 * 0ml] + [0.5 * 0.5ml]). The first step of the procedure involved presenting the monkeys with choices between this gamble and varying safe rewards (in 0.05 ml increments), from these, the safe reward that was equivalent to the gamble in utility terms was identified (i.e. the safe reward chosen in equal proportion to the gamble; see Fig. 1c).

To estimate this safe reward, the following logistic sigmoid curve was fitted to the proportion of safe choices for each of the gamble/safe pairing:

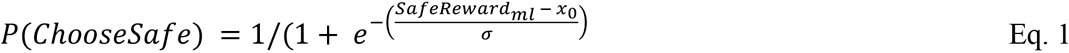

Where probability that the monkeys would choose a safe reward over the 0.5 utility gamble (P(*ChooseSafe*)) was contingent on the safe option’s magnitude (*SafeReward_ml_*) and two free parameters: x_0_, the x-axis position of the curve’s inflection point, and σ, the function’s temperature. Importantly, this function’s inflection point represented the exact safe magnitude for which the monkeys should be indifferent between the set gamble and a given safe reward. The x_0_-parameter could thus be used as a direct estimate of the gamble’s certainty equivalent (CE), or, put simply, the safe reward equivalent to a utility of 0.5. Only sequences that contained a minimum of three different choice pairs (repeated at least 4 times) were used in the elicitation of CEs.

From the CE identified as the 0.5 utility value, two new equiprobable gambles were created representing utility values of 0.25 (¼ of the utility range) and 0.75 (1/4 and ¾ of the utility range, respectively). Of the two new gambles, one was set between 0 ml and the first CE’s ml value, the other was set between the first CE and 0.5 ml (Fig. 2b). The CE elicitation procedure (logistic fitting, Fig. 1c) was repeated for each of these gambles. Crucially, gamble/safe pairings for both gambles were interwoven in the same sequence – to ensure a similar spread in the presented rewards.

After eliciting the CEs of these gambles, the estimation procedure was repeated one final time with the new CEs as the upper or lower gamble outcomes. Here, the fractile procedure would automatically terminate if no safe rewards could fit between the outcomes of the new gambles; this would occur if the animal was particularly risk-seeking or risk-averse. If this was the case, utilities of 0.25, 0.5, and 0.75 would be mapped onto the appropriate reward magnitudes and the elicitation sequence would end. If, instead, the three fractile steps were successful, the procedure would result in a mapping of five utilities, 0.125, 0.25, 0.5, 0.75, and 0.875, onto five safe rewards. Only sequences where at least 3 utility points were successfully identified were used in the study (monkey A: 22 sessions; monkey B: 7 sessions).

### Estimating utility functions in riskless choice

For riskless choice sequence, choice ratios between pairs of safe options were measured-this time looking at the likelihood of a monkey choosing the high magnitude option over the lower magnitude one (Fig. 1d). The range of juice rewards (0.05 ml to 0.5 ml) was divided into sets of 0.05 ml increments and safe-safe pairs centered on these magnitude increments. For each increment, we defined three sets of safe-safe choices where each pairing differed by 0.02 ml, 0.04 ml, or 0.06 ml. The small size of these differences ensured that choices would be stochastic. These differences are hereafter defined as ‘gaps’, i.e. safe-safe pairings of fixed differences, where three sets of gaps were anchored at each incremental ‘midpoint’.

The likelihood of choosing the higher magnitude option in different gap-midpoint pairings was used to infer the shape of the monkeys’ utility functions (Fig. 3a, b, c). Specifically, the difference between the likelihoods of choosing the better options, at different midpoints, reflected the separability of the utility of different reward magnitudes. Under RUM, the degree of certainty with which choices are made (i.e. the closer choice ratios are to 100%) directly correlates with the separability of the noisy utilities that correspond to each option in a choice. This implies that, looking at repeated choices between two set magnitudes, a decision-maker with a flatter utility function should exhibit more stochasticity in their choices (i.e. less precision) than a decision-maker with a steeper utility (i.e. more precision). Changes in choice ratios between sequential midpoints, as averaged across gaps, could therefore be used as a proxy for a monkeys’ utility slope.

**Figure 3.**
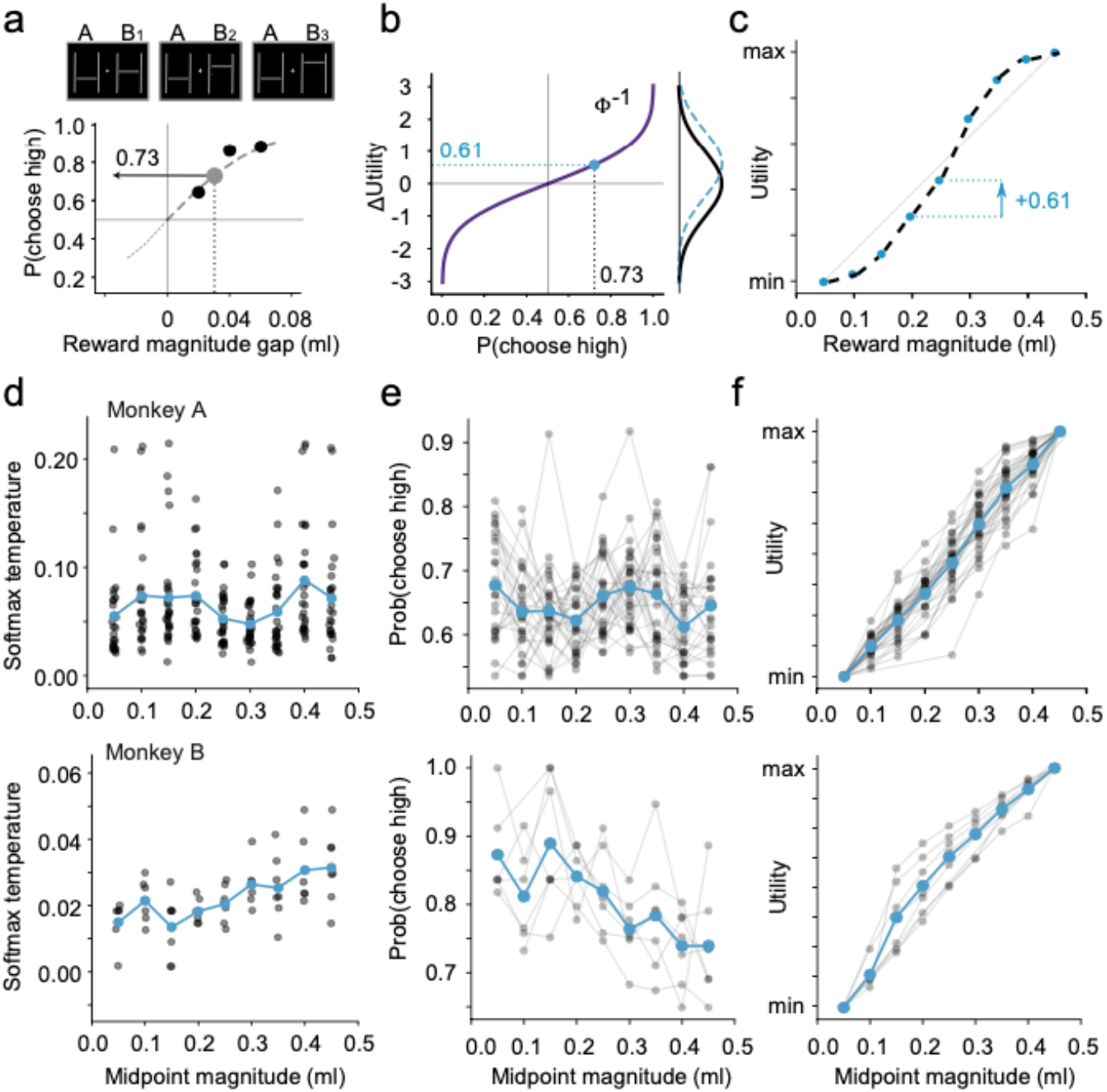
Estimating riskless utilities from the stochasticity in safe-safe choices. a) **Measuring stochasticity in choices between safe reward pairs**. Example visual stimuli (top) representing choices between safe rewards (A: low, B: high) resulting in different percentage of choices for the high option (bottom; black dots). This was repeated for different rewards pairs, centered at different increments (midpoints). For each midpoint, the likelihoods were fitted with a softmax curve (dashed), used to estimate the probability of choosing the larger option for a gap of 0.03 ml (gray dot). b) **Choice ratios as differences in utility**. The likelihoods that monkeys would pick the better reward were transformed using the inverse cumulative distribution function (iCDF) of a logistic distribution. The utility of different rewards took the form of equally noisy distributions centered at the monkeys’ ‘true’ utilities. The output of iCDFs is the distance between these random utilities (i.e. the marginal utility). c) **From marginal utilities to utility**. The cumulative sum of marginal utilities approximated a direct utility measure for each midpoint. These measurements were normalized whereby the utility of the highest midpoint was 1, and the starting midpoint had a utility of 0. d) **Daily strength of preference estimates**. Each point represented the temperature of the softmax curve fitted on the choice ratios (blue points: average across days). The lower the temperature parameter, the steeper was the softmax curve and the more separable were the random utilities. Lower values meant higher marginal utility measurement (steeper utility function), higher ones meant lower marginal utility (flatter function). e) **Daily choice ratio estimates from softmax fits**. Estimates from the same day are linked by grey lines. Ratios of 0.5 meant that the random utility of the two options were fully overlapping (i.e. flat utility function); choice ratios closer to 1 meant random utilities that were fully dissociated and non-overlapping. f) **Utility functions**. Utilities estimated in single days (grey lines) and averages (blue), normalized relative to the minimum and maximum midpoint.

To estimate these RUM-compliant utilities, logistic curves were fitted to the likelihood of choosing the better option (for the three gaps) at every midpoint level (Fig. 3a)

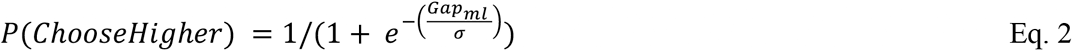

Unlike for CE estimation, this logistic function captured the likelihood of choosing the high-magnitude option (in a safe-safe pairing) contingent on the gap between the two options (*Gap_ml_*) and σ, the logistic function’s temperature. Just as is the case for CE estimation however, the utility estimates relied on aggregate choices between multiple reward pairs. The logistic fit also highlighted sequences where monkeys would not follow even the most basic principle of rational choice: weak stochastic dominance (picking an objectively lower outcome). Choices where this was the case were removed from all future analyses: that is, when the estimated temperature parameters of logistic fits were negative (i.e. the larger the gap, the lower the likelihood of choosing the better option) or significant outlier (p <0.05; Grubbs’s test). In monkey A, 38 choice sets were removed from a total of 279 choice sets (14 negative parameters and 24 outliers). In monkey B, 1 choice set was removed from a total of 62 choice sets (1 negative parameters and no outliers).

Where logistic fittings were successful, the functions were used to estimate the higher-lower choice ratio, at each midpoint, for an untested magnitude gap of 0.03 ml (Fig. 3a). Then, the inverse cumulative of a logistic probability density function (centered at 0 with variance = 1) was used to estimate the distance, in utility terms, between the two magnitudes in the 0.03 ml gap (Fig. 3b). In other words, these 0.03 ml gaps were placed onto a shared scale (i.e. random utilities) through the assumption that, on each trial, the probability that the monkeys would pick the better reward (*x_i_*) was given by:

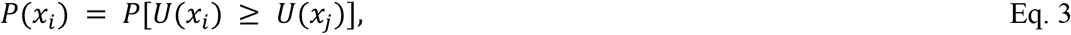

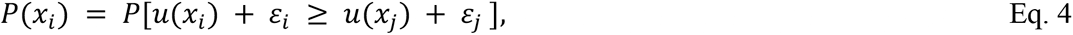

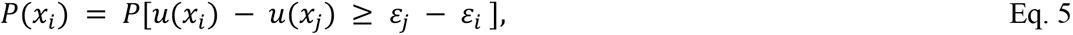

In this form, the probability of choosing ***x_i_*** rather than ***x_j_*** was given by the probability that the difference in the true utilities of ***x_i_*** and ***x_j_*** was greater or equal to the noise on ***x_j_***(***ɛ_j_***) minus the noise on ***x_i_***(***ɛ_i_***). From this, it followed that the distribution of noise differences could be used as a predictor of the distance between the two true utilities (***u***(***x*_*i*_**) and ***u***(***x*_*j*_**)). Because of the assumption of constant noise, the probability of choosing ***x*_*i*_** over ***x*_*j*_** would be directly proportional to the distance between the true utility of two options. In accordance with McFadden’s formulation (McFadden, 1974, 2005; Stott, 2006), we assumed that the distribution of error differences (*ε_j_* − *ε_j_*) took a logistic form:

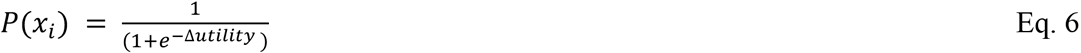

and then used the inverse of this logistic distribution’s CDF to estimate the difference in utilities (Δ*utility*) between the hypothetical 0.03 ml reward gaps (Fig. 3c)-essentially the slope of the monkeys’ utility function at every midpoint. The cumulative sum of these slopes provided an estimate of the utility at each midpoint.

### Modelling risky and riskless choices in a common metric

Because the utilities measured from aggregate behavior did not account for probability weighting on choices (i.e. they were EUT utilities rather than PT ones), parametric utility functions were re-estimated from individual choices using a discrete choice model that could account for the effects of both, separately. This placed utility metrics for risky and riskless choices on a common and comparable scale, and, importantly, it allowed for the inclusion of probability weighting as an additional contributor to the monkeys’ preferences.

As in most discrete choice models (and in line with the aggregate RUM metric), a logit function (softmax) was used to represent noise in the decision-making process. The probability of the monkey making either a left or right choice was therefore given by:

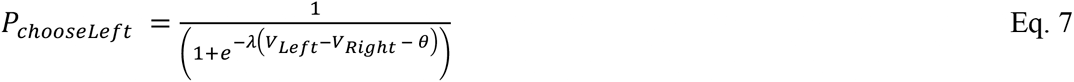

Where the probability of choosing the left option is a function of the difference in value between the left and right options, the noise parameter, *λ*, and the side bias parameter *θ*. The value of each option (V_Left_, V_Right_) took on the functional form prescribed by PT in its cumulative form (Tversky & Kahneman, 1992):

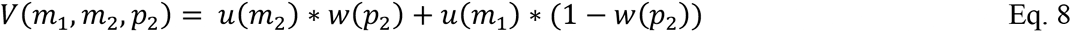

where *m_1_* and *m_2_* were the low and high outcome magnitudes respectively, while *p_2_* was the probability of obtaining the high outcome; the probability weighting function (*w(p)*) corresponded to a power function:

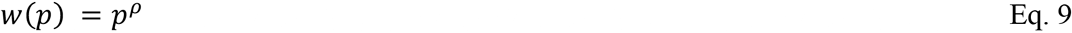

The utility of the option’s outcome (*u(m)*) was the CDF of a two-sided power distribution (Kotz and Dorp, 2010):

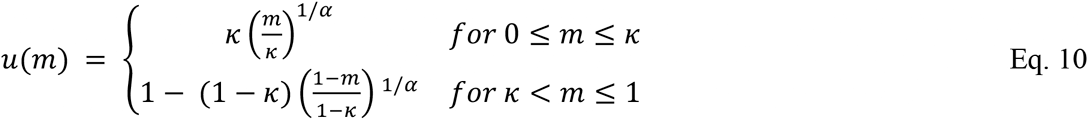

In the probability weighting function, the ρ-parameter prescribed either an overweighing (ρ>1) or underweighting (ρ <1) of an outcome’s probability. The utility measure was a function of an α-parameter and an inflection point κ, where the curvature of the utility function would invert. Each outcome magnitude (*m*) was normalized onto a 0-1 scale, so that κ was bounded by the range of outcome magnitudes experienced by the monkeys (values from 0 to 1, corresponding to 0 ml and 0.5 ml respectively).

Each of these parameters was fit to single-choice data by maximizing the sum of 21 likelihoods defined on the model as

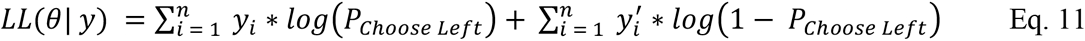

For each individual choice trial (*i*), *y* and *y’* indicated a left or right choice respectively (1 if yes, 0 if no), n was the total number of trials for the session, and P_*Choose Left*_ was the output of the earlier logistic function (Eq. 7). This discrete choice analysis was restricted to choice sequences previously deemed appropriate for the aggregate preference estimations described in earlier sections.

### Statistical comparison of risky and riskless choices

Estimating utilities through discrete choice modelling allowed for the comparison of the functional parameters that best described the monkeys’ decisions in risky and riskless choices, and to explore the unique contributions of both magnitudes (through utility) and probabilities (through probability weighting) in a way that aggregate, non-parametric measures did not permit.

Because the logit function’s *λ*-, and the utility’s α-parameters were asymmetrically distributed (with positive values <1 accounting for as much change as values >1), these were log-transformed before proceeding with any comparison. Then, the parameters elicited in risky choice sequences were compared to those estimated from riskless sequences using a one-way multivariate analysis of variance (or MANOVA) whereby the main comparison factor in the analysis was the risk-riskless choice scenario described by each set of parameters. Since the probability weighting parameter for riskless choices was constant and fixed at 1, we restricted the MANOVA analysis to the softmax and utility parameters. We then ran additional correlation analyses (Pearson’s R) between risky and riskless utility parameters to determine if the parameters in one set of choices could predict those of another.

All parameters were compared independently for each monkey, results were never pooled across animals, and the statistics for each monkey are reported separately. All statistical analyses were considered significant at p < 0.05.

## Results

### Experimental design

Prospect theory implicitly assumes that the utilities that guide risky and riskless decisions are the same. We sought to validate this assumption in macaque monkeys by comparing the decisions they made in risky versus riskless choices. Two rhesus macaques were trained to make choices between pairs of reward options presented on the left and right sides of a computer screen by moving a joystick towards the chosen side (Fig. 1a). The reward options varied in terms of blackcurrant juice quantity as well as in the probability that they would be delivered. The monkeys received the selected rewards after every trial – contingent on their delivery probability.

Choice preferences were elicited in trial sequences in which either both options were certain and therefore riskless, or in sequences where one option was certain (safe option) and the other was a risky gamble with two possible outcomes (juice magnitudes), each delivered with probability p = 0.5 (equiprobable gamble). We separately used these riskless or risky choices to infer an animal’s utility function, compatible with PT. Choice sequences were structured in a way that allowed us to map utilities onto aggregate behavioral metrics, and to then model these choices under the assumptions of Prospect Theory.

In risky choices, utilities were estimated by psychometrically measuring the certainty equivalent (CE) of equiprobable gambles and then applying the fractile method, a stepwise procedure whereby one progressively sections the range of possible rewards using the CEs estimated from previous steps (see Methods section). In each session, we obtained five intermediate points of the EUT-compatible utility functions (Fig. 2).

Since gambles were off-limits to estimate riskless utilities, the random utility maximization (RUM) framework was used in riskless choices to estimate utility differences between two reward magnitudes (Fig. 3a, b). The utility functions were then reconstructed by cumulatively summing all such utility increments (Fig. 3c). This procedure produced seven utility levels, corresponding to our discrete estimate of the RUM-compatible utility function (see Methods section) (Fig. 3).

### Utility functions in risky and riskless choice

Choice measurements from risky and riskless sequences were gathered on the same day, in 22 and 7 sessions for monkeys A and B respectively. We used these choices to estimate the utility function underlying the measured choice pattern.

For both risky and riskless sequences, a link between utility measurements and reward magnitudes was confirmed via one-way ANOVA. Both monkeys exhibited a significant main effect of utility on the CEs (Fig. 2c) in risky choices (Monkey A: F_(4,124)_ = 35.482, p = 9.763∙10^−20^, Monkey B: F_(4,39)_ = 172.537, p = 3.090∙10^−24^). In riskless choices, we contrasted the utilities with the midpoint reward magnitude (Fig. 3f), highlighting a significant main effect (Monkey A: F(8,232) = 375.763, p = 3.503∙10^−128^; Monkey B: F(8,52) = 85.561, p = 3.474∙10^−27^). These basic results illustrated how the utilities associated with different reward magnitudes were significantly different from each other, which would not have been the case if monkeys selected options at random.

Importantly, the utility levels were significantly rank-ordered in relation to the reward magnitudes (Spearman rank correlation in Monkey A: risky Rho= 0.7209, p=5.853∙10^−22^; riskless Rho= 0.9628, p=8.035∙10^−138^. In Monkey B: risky Rho= 0.9446, p=6.092∙10^−22^; riskless Rho= 0.9665, p=1.529∙10^−36^), in line with the fundamental principle of utility functions being monotonically related to the reward magnitudes. In general, utilities appeared to be non-linear functions of physical reward magnitudes.

In risky choices, the full elicited risky utility functions followed an S-shape pattern in both monkeys, reflecting the typical risk attitudes observed in macaques: risk-seeking (convex utility) for relatively low-magnitude rewards and risk-aversion (concave utility) for relatively high-magnitude ones (Fig. 2c).

In riskless choices, we compared the estimated utility increments in order to highlight any non-linearity in the utility shape. As increments in utility were proportional to the temperature parameter (i.e. the slope) of the softmax curves that described choices around a certain magnitude level, the softmax temperature could be used as a proxy for linearity: a constant temperature across magnitude levels would correspond to a linear utility function, while a varying temperature would indicate non-linear utility. We compared the temperature parameter across midpoints and found that it varied significantly with magnitudes (Fig. 3d; Monkey A: F_(8, 232)_ = 2.663, p = 8.165∙10^−3^); Monkey B: F_(8, 52)_ = 4.187, p = 6.370∙10^−4^) highlighting the non-linearity in the riskless utility function, in both monkeys. The softmax temperature, as a function of the midpoint, reached a minimum (around 0.30 and 0.15 ml for monkeys A and B respectively) before increasing again, suggesting a slight S-shape for the riskless utility function (Fig. 3f).

Although these aggregate utility measures were based on commonly defined economic models, they were not (i) PT-compatible, and (ii) comparable between the risky/riskless choice scenarios. In fact, we estimated the risky utility functions following EUT, which, in contrast with PT, assumes no subjective weighting of probabilities; the utility functions had different magnitude-ranges in risky and riskless choices (0 to 0.5 ml and 0.05 ml to 0.45 ml, respectively) and different discrete steps. We sought to overcome these limitations by defining a utility estimation method that allowed for a direct comparison of utility in risky and riskless choices, compatibly with economic choice models.

### Risky and riskless utility functions on a common scale

To directly compare the utility functions between risky and riskless choices, we re-estimated utilities on a common scale, compatible with PT. We used the same discrete choice model (Eq. 7) to describe both riskless and risky choices, without the need of two different estimation procedures.

The main assumption of our model is that a random quantity is added to each option’s utility at every trial, using the PT model as the underlying deterministic choice mechanism. This model introduced stochasticity in choices and could readily be applied to both risky and riskless choices without modification.

In the model, utility functions took the form of the cumulative distribution function of a two-sided power distribution (Eq. 10; Kotz & Dorp, 2010), a 2-parameter function that could easily account for complex risk-attitudes (Kontek and Lewandowski, 2018): if *α* < 1, the utility function would be convex and predict risk-seeking choices up to the inflection at parameter *κ* (predicting risk-averse choices thereafter); if instead *α* > 1, the utility function would be concave and predict risk-averse behavior up to the inflection at *κ* (predicting risk-seeking behavior afterwards). For risky choices, a 1-parameter power function captured the weighting of probabilities (Eq. 9). Since the only probability experienced was p = 0.5, *ρ* > 1 implied an overweighting of the probability of receiving the highest reward whilst *ρ* < 1 implied underweighting.

We defined three forms of this discrete choice model, with different free parameters: the EV model (linear utility and probability weighting), where only the “noise” parameter was free to vary; the EUT model (linear probability weighting), where the utility parameter could vary; and the PT model, with both utility and probability weighting free parameters. In risky choices, we compared the goodness-of-fit of the three models to identify the one that would produce the best estimate of a utility function. In riskless choices, we estimated the utility function using the EUT model.

In risky choices (Fig. 4a, b), both the EUT and PT models predicted s-shaped utility functions (Monkey A EUT: t(22)= −29.0190, p<0.00001; Monkey A PT: t(22)= −28.2543, p<0.00001; Monkey B EUT: t(22)= −4.2859, p=0.005172; Monkey B PT: t(7)= −7.4532, p=0.000301). The PT model, however, relied on concave probability weighting (one-sample t test, Monkey A: t_(22)_ = − 4.2533, p = 3.55∙10 ^−4^; Monkey B: t_(7)_ = −2.7316, p = 0.0341), rather than a convex utility function, to explain risk-seeking behavior. For that reason, PT’s s-shaped utility functions were mostly left-skewed (more concave than convex) whereas EUT utility functions captured risk-seeking behavior solely through a right-skewed s-shape (more convex than concave) (Fig. 4a). Overall, the daily best-fitting parameters from the PT and EUT models were significantly different from each other (Table 1), with the PT model capturing behavior significantly more reliably than both EV and EUT models (Fig. 4b; Wilcoxon rank sum test; monkey A: p = 1.0∙10^−4^; monkey B: p = 1.8∙10^−2^). Through the PT model, we could separate the contribution of utility and probability weighting to the risk attitude, obtaining a better estimate of the utility function underlying choices, compared to the EUT model.

**Figure 4.**
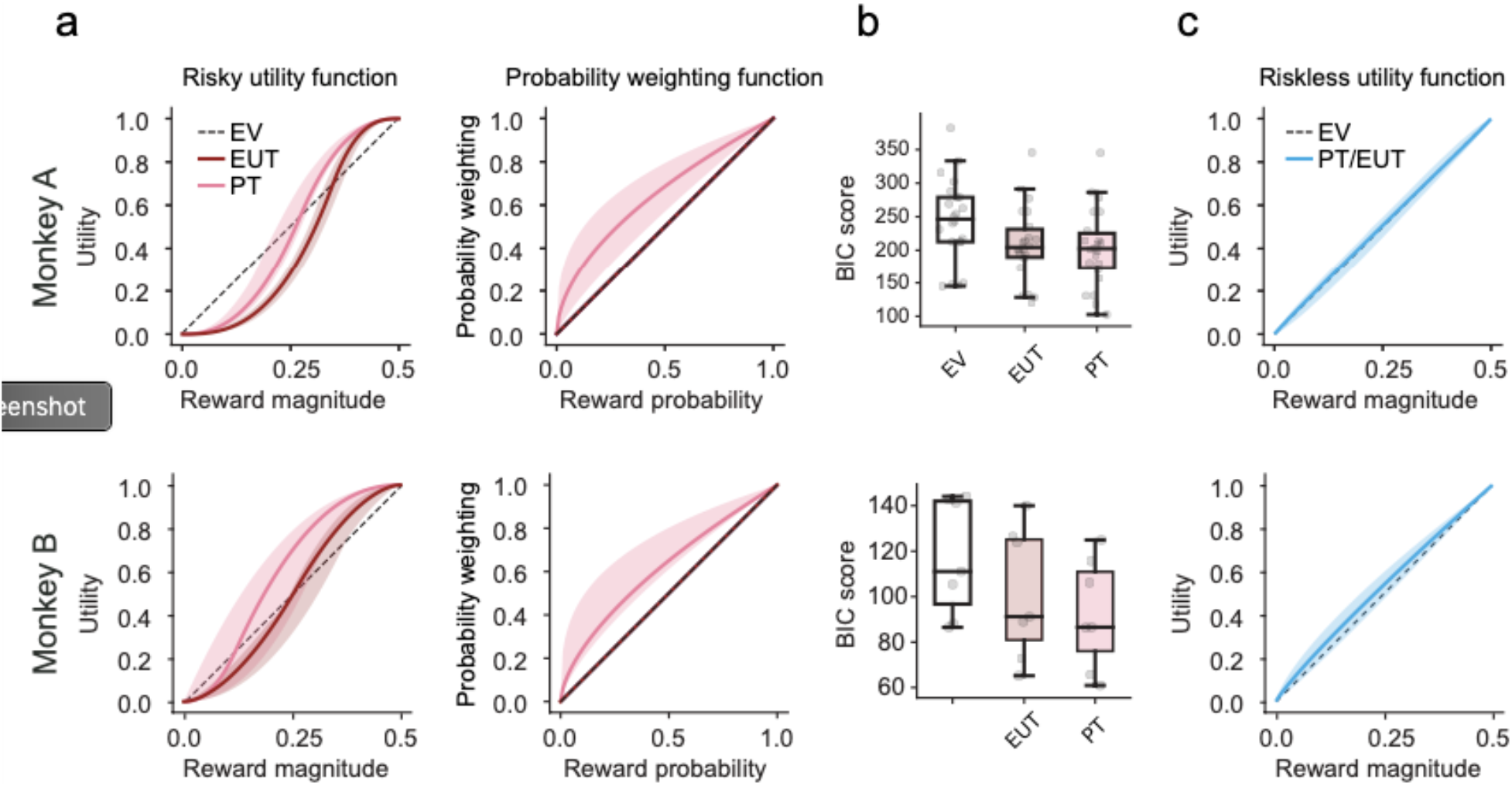
Discrete choice estimates differ between risky and riskless choices. a) **Utility functions in risky choice.** Median parametric estimates for utility functions and probability weighting functions fitted to risky choices. Shaded area: 95% C.I. on the median of these functions. Two versions of the discrete choice model were fitted: the expected utility theory (EUT) model predicted choices solely based on reward options’ utilities (without probability weighting); the prospect theory (PT) model, predicted choices based on utilities and probability weighting. An expected value (EV) based model was included for comparison. Monkeys were risk-seeking, but where the PT model accounted for this mainly through probability weighting, the EUT model accounted for it through a more convex utility. b) **Comparison of risky choice models**. The PT model described individual choices better than EUT and EV. Bayesian information criterions (BIC) were calculated from the log likelihoods of the daily best-fitting PT and EUT discrete choice models. c) **Utility functions in riskless choice**. Median parametric estimates for utility functions fitted to riskless choices (shaded area: 95% C.I. on the median). The discrete choice model predicted choices from the expected utilities of rewards (no probability weighting). Utilities were mostly linear, though slightly concave.

**Table 1.** MANOVA Tests for pairwise differences between the risky EUT, risky PT, and riskless discrete choice models.

| | Utility Type | F (1, 42) | p | Wilks $\lambda$ |
| --- | --- | --- | --- | --- |
| Monkey A | Riskless,<br>Risky (PT) | 28.697 | $6.158 \cdot 10^{-12}$ | 0.209 |
| | Risky (EUT),<br>Risky (PT) | 5.475 | $6.856 \cdot 10^{-4}$ | 0.581 |
| | Utility Type | F (1, 12) | p | Wilks $\lambda$ |
| Monkey B | Riskless,<br>Risky (PT) | 8.744 | $4.239 \cdot 10^{-3}$ | 0.155 |
|  | Risky (EUT),<br>Risky (PT) | 1.687 | 0.243 | 0.487 |

The analyses were run on four of the five free parameters, excluding probability weighting. The risky EUT and riskless models had no probability weighting parameter to compare with the risky PT model's probability weighting.

**Table 2.** Two-way ANOVA Tests for pairwise differences between three sets of certainty discrete equivalents.

|  | Utility Type | Df | F (1, 60) | p |
| --- | --- | --- | --- | --- |
| Monkey A | Risky/Riskless,<br>Risky (PT) | (7, 168) | 432.024 | $6.859 \cdot 10^{-104}$ |
| | Risky/Riskless,<br>True CEs | (7, 142) | 118.972 | $3.665 \cdot 10^{-56}$ |
| | Risky (PT),<br>True CEs | (7, 142) | 98.785 | $2.3 \cdot 10^{-51}$ |
|  | CE Type | Df | F (1, 60) | p |
| Monkey B | Risky/Riskless,<br>Risky (PT) | (9, 60) | 193.653 | $6.75 \cdot 10^{-41}$ |
| | Risky/Riskless,<br>True CEs | (9, 149) | 208.550 | $9.462 \cdot 10^{-80}$ |
| | Risky (PT),<br>True CEs | (9, 149) | 211.873 | $3.189 \cdot 10^{-80}$ |

The certainty equivalents were derived from the daily predictions of the risky/riskless hybrid model, the PT model, and the ones measured from out-of-sample sequences.

In riskless choices (Fig. 4c), the utility function’s α parameter was not significantly different from one (t test, Monkey A: t(22)=−0.3267, p=0.7471; Monkey B: t(7)=1.3457, p=0.2270). This implied that the riskless utility functions were close to linear, suggesting that magnitudes were objectively represented, according to the RUM framework.

### Mismatch between risky and riskless utility functions

When comparing riskless and risky utilities computed on a common scale, we found a significant difference in the utility functions’ shapes in terms of α parameter, in both monkeys (Fig. 5a; Monkey A: F_(1,42)_ = 72.717, p = 1.04∙10^−10^; Monkey B: F_(1,12)_ = 24.221, p = 3.52∙10^−4^).

**Figure 5.**
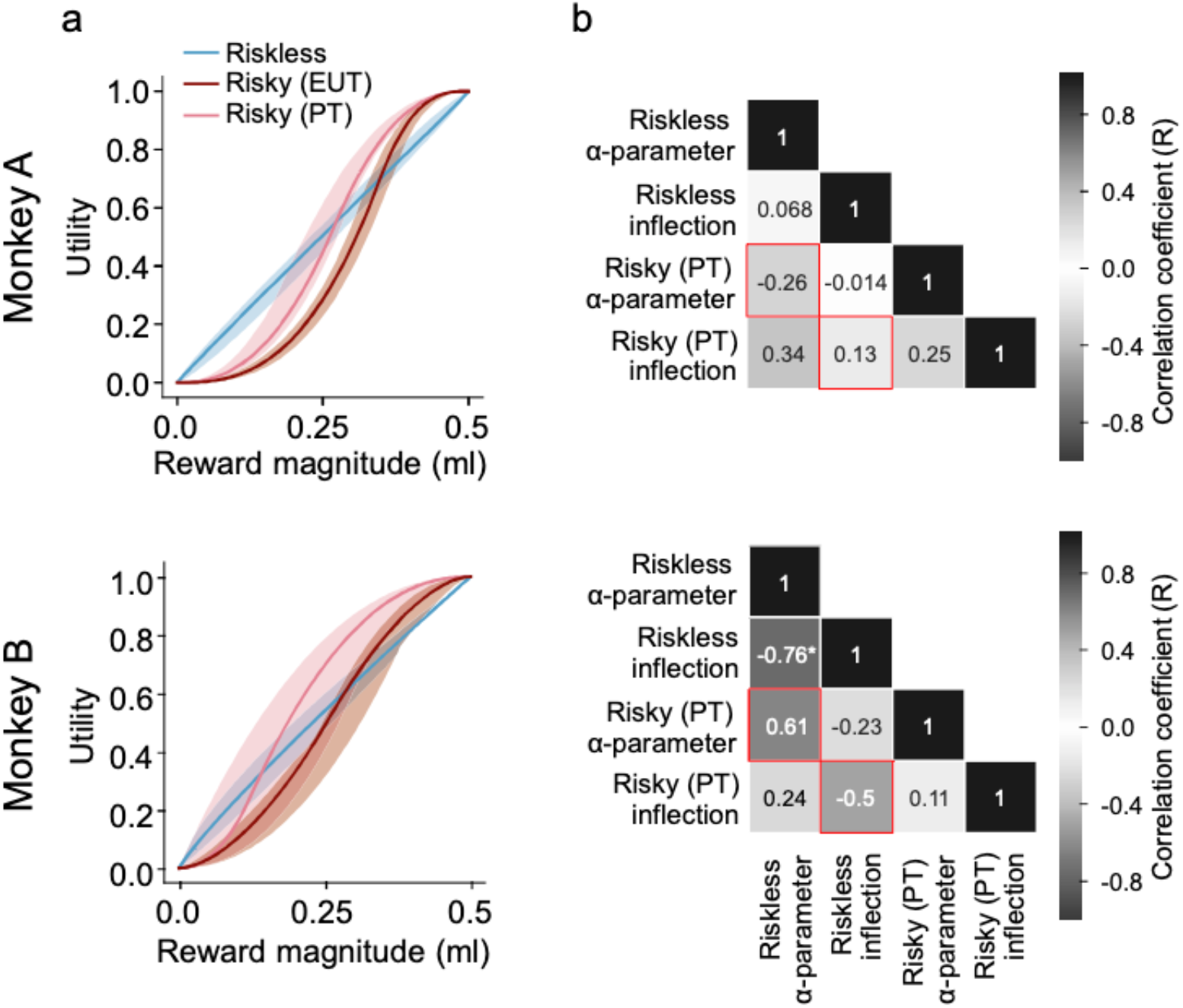
Risky utilities do not predict riskless ones, and vice-versa. a) **Median utility function estimates for risky and riskless choices.** The shaded area represents the 95% C.I. on the median of these functions. For riskless choices, utility estimates were mostly linear (though slightly concave). For risky utilities, the two different versions of the discrete choice model predicted S-shaped utilities, but risky EUT utility functions were more convex than PT utility functions. b) **Absence of correlation for utility parameters in risky vs. riskless choices**. Pearson’s correlations were run on the parameters from risky and riskless scenarios. Red squares highlight Pearson’s R for the correlation of the α and inflection parameters between risky and riskless choices. Asterisks (*) indicate significant correlations (p < 0.05).

Monkey B’s difference in the utility’s inflection point between risky and riskless choices (Monkey A: F_(1,42)_ = 1.282, p = 0.264; Monkey B: F_(1,12)_ = 17.153, p = 0.00136) was significant, while we found no significant difference in either the noise or the side bias parameters (noise: Monkey A: F_(1,42)_ = 2.760, p = 0.104; Monkey B: F_(1,12)_ = 0.182, p = 0.677; side bias: Monkey A: F_(1,42)_ = 0.2407, p = 0.626; Monkey B: F_(1,12)_ = 2.338, p = 0.152).

Overall, these results show that the dissimilarity between the modeled riskless and risky choices was mainly due to a difference in the non-linearity of the utility functions, as expressed by the α parameter. The utility function was strongly non-linear in risky choices, while it was close to linear in riskless choices.

The difference in utility functions was also evident when comparing risky and riskless data from single days, through a correlation analysis: we found was no significant correlation between any of the parameters of risky utility functions and those of riskless utility functions across days (Fig. 5b).

As a control, we correlated the measured riskless choice percentages (for the hypothetical 0.03 ml gap, grey dot in Fig. 3a) with the modeled ones, separately using the utility function elicited from risky or riskless choices. We found a significant correlation coefficient when predicting riskless choices using the riskless utility function (Monkey A: R=0.442, p=1.278∙10^−9^ Monkey B: R=0.484, p=7.610∙10^−5^) but not using the risky one (Monkey A: R=0.104, p=0.175 Monkey B: R=0.087, p=0.503). This confirmed that the riskless utility function captured the behavior in riskless choices while the risky utility function did not, emphasizing the difference in risky/riskless utilities.

In summary, estimating utilities through PT, rather than EUT, brought risky fits more in line with riskless ones (Table. 1; Fig. 5a) in line with previous human studies (Stalmeier and Bezembinder, 1999; Abdellaoui et al., 2007). However, a direct comparison between the risky and riskless utility parameters revealed significant differences in the utility functions’ shapes between the two choices scenarios (Fig. 5b).

## Discussion

Using a robust, incentive-compatible task, we showed that utility functions that describe decisions involving risk more closely mimicked riskless utility functions, if probability weighting was considered. We modelled macaque monkeys’ risky and riskless choices through stochastic versions of PT and EUT, and reliably estimated functional parameters that best described their choices. Each day, the monkeys were presented with risky or riskless binary choice sequences. In risky ones, they made choices between gambles and safe rewards; in riskless ones, both choices had a single, certain outcome. We found that modelling monkeys’ risky decisions via the PT model of choice, in addition to providing a better fit than EUT, led to decision parameters that more closely resembled riskless ones. This trend is in-line with the human literature (Stalmeier and Bezembinder, 1999; Abdellaoui et al., 2007). However, the direct comparison differed: the monkeys’ utility functions elicited in riskless and risky choices were more alike, but they were still significantly different.

In terms of behavioral metrics, the CEs estimated in fractile sequences suggested both monkeys were risk-seeking for all but the highest of reward magnitudes that they experienced. The PT and EUT models predicted similar risk-seeking behavior via an overweighing of gamble options, but they differed in the way in which they achieved this. Both EUT and PT models predicted s-shaped utilities, the PT model, however, accounted for the monkey’s risk seeking behavior mostly through its concave probability weighting. In other words, the subjective probability of ‘winning’ a gamble was higher than the objective probability of winning regardless of utility’s effects. EUT fits, on the other hand, captured risk-seeking behavior exclusively through their utility function; one that was right-leaning (more convex than concave) and so predicted higher utilities for gamble options than for safe ones. Since PT’s utilities were ‘free’ from the effects of probability weighting, the s-curves were left-shifted (i.e. more concave than convex), suggesting a relatively more risk-averse utility function than from EUT’s predictions. Comparing these findings to the riskless utility fits, we found that PT utilities deviated far less from riskless utilities than EUT ones. Still, the utilities estimated from riskless binary choices were relatively linear (if slightly risk-averse), a shape that was at odds with that of the risky PT estimates. It appears that, at least within the confines of our experiment, the difference between risky and riskless utilities was not as simple as the addition of a probability weighting parameter.

Assuming that the discrete choice model is correct, the difference in utility functions for risky and riskless utilities could be used as a quantitative basis for a neuronal test of utility coding. By recording the activity of single neurons during risky or riskless choices, the pattern of neuronal activations in utility-coding neurons should reflect the different utility shapes elicited though behavior in the two choice scenarios.

As an alternative interpretation, the source of discrepancy between risky and riskless utility function could be due to limits in the model specification. Alternative models should be compared to support this hypothesis, including different assumptions on the noise shape: while the current model assumes a constant and symmetric noise around each option’s utility, this could be an oversimplification. A more biologically plausible contribution of noise on the utility measure could include asymmetric and non-constant noise (especially for activity rates close to the limits of the neurons’ dynamic range) as well as noise applied separately to every option’s component (magnitudes and probabilities).

Moreover, monkeys could be using different strategies for solving the risky and riskless choice problems, implying different brain mechanisms. In particular, riskless choices closely resemble a perceptual discrimination task, in which subjective values would not be required and the optimal solution would be to perceptually compare the visual stimuli.

While the same binary choice design was used in risky and riskless choices, the difference between options was much greater in risky sequences than in riskless ones. To estimate aggregate riskless utilities, for example, the rewards that the monkeys experienced differed only by up to 0.06ml in every trial. In risky sequences, on the other hand, gambles were pitted against safe rewards spread over the full range of the gambles’ outcomes. Monkeys experienced a broad range of magnitudes in each of the sequences, but the differences between riskless choices could have required far more attention to dissociate than those in riskless choices (something we cannot account for; but see, Farashahi et al., 2018).

Where these findings fail to replicate the data from risky and riskless introspective studies (though see Hertwig, Wulff, & Mata, 2018), they are nonetheless in line with the incentive-compatible time trade-off approach. Since these types of time discounting tasks are easily adapted to study preferences in rhesus macaques (Hayden and Platt, 2007; Kobayashi and Schultz, 2008; Hwang et al., 2009; Blanchard et al., 2013), it would be interesting to see how utility functions estimated using time trade-offs in macaque monkeys correlate with the present findings. Another approach that would be interesting to consider is the one used by Chung, Glimcher and Tymula (Chung et al., 2019), where they compared risky and riskless choices between bundles of outcomes-estimating utilities through identifying the combinations of rewards for which decision-makers are indifferent. They found that risky and riskless choices could be reconciled when choices involved gains, but that PT failed to reconcile the two when the choices involved losses. Since preferences over losses are generally risk-seeking (for humans), it could be that the macaque monkeys’ risk-seeking behavior mimics this loss-related discrepancy. If macaque monkeys were to, in risky settings, adjust their expectations in a way that paints the lower outcome of a gamble as a loss, one would expect the lower end of their utility function to behave like the loss side of PT’s value function (Kahneman and Tversky, 1979). There is some evidence that rhesus macaques (and indeed humans) do this: they exhibit preferences consistent with win-stay lose-shift strategies (Gilovich et al., 1985; Barron and Erev, 2003; Heilbronner and Hayden, 2013). For repeated gamble-safe choices, they generally reverse their risk-seeking preferences for gambles depending on if they have previously won or lost a previous gamble instance (Lau and Glimcher, 2005; Blanchard et al., 2014; Ferrari-Toniolo et al., 2019b). If this is the case, fitting macaques’ choices through utility models that account for trial-by-trial changes in preference functions are likely to do a better job at reconciling risky and riskless utilities than using fixed utility and probability weighting functions applied to the entire experimental procedure.

Overall, the results presented here add to the need for decision models to account for flexible, context-specific preferences (Hayden, B; Heilbronner, S; Nair, A; Platt, 2013; Heilbronner and Hayden, 2016; Farashahi et al., 2018). For decision-theory as a whole, reconciling dynamic preferences with more traditional economic models would go a long way to making more accurate, descriptive predictions.

## Acknowledgements

We thank Aled David and Christina Thompson for animal and technical support. The work was funded by the Wellcome Trust (WT 095495, WT 204811) and the ERC (Advanced Grant 293549).

## Notes

Conflict of Interest: The authors declare no competing financial interests.

### Competing Interest Statement

The authors have declared no competing interest.

